# An ensemble of *Gleditsia sinensis Lam*. and gut microbiota against alcoholic liver disease

**DOI:** 10.1101/2023.05.31.543000

**Authors:** Ki-Kwang Oh, Sang-Jun Yoon, Su-Been Lee, Sang Youn Lee, Haripriya Gupta, Raja Ganesan, Satya Priya Sharma, Sung-Min Won, Jin-Ju Jeong, Dong Joon Kim, Ki-Tae Suk

**Affiliations:** Institute for Liver and Digestive Diseases, College of Medicine, Hallym University, Chuncheon 24252, Korea

**Author notes:** Correspondence, Orcid ID: 0000-0002-9206-9245.

**Keywords:** *Gleditsia sinensis Lam.*, gut microbiota, equol, Bauer-7-en-3-one, Urs-12-en-3-one

## Abstract

*Gleditsia sinensis Lam.* (GSL) is a medicinal herb and a noticeable resource of possessing hepatic protective agents such as alcoholic liver disease (ALD). At present, it has been documented that gut microbiota (GM) is related directly to etiology of ALD. Nevertheless, the bioactive molecules in GSL, favorable GM, targets, and key mechanism(s) against ALD are yet to be revealed. Hence, we integrated the significant four components to clarify the nuanced pathogenesis with help of network pharmacology (NP) concept. We retrieved significant metabolites via gutMGene and constructed GSL or GM-Signaling pathways-Targets-Metabolites (GGSTM) networks. Finally, molecular docking test (MDT) was performed to verify the key findings. The gutMGene suggested that 16 GM and 6 metabolites were related to the two signaling pathways through GGSTM networks. Both MDT and frontier molecular orbitals (FMO) theory revealed the most stable conformers: equol from *Lactobacillus paracasei JS1* on IL6, Bauer-7-en-3-one, and Urs-12-en-3-one from GSL on PPARA, PPARD, and PPARG, respectively. In conclusion, this study sheds light on the combinatorial effects of GM, and GSL in treating ALD via systems biology concept.

## Introduction

Alcoholic liver disease (ALD) includes diverse hepatic-related disorders: hepatic steatosis (HS), hepatic cirrhosis (HC), even hepatocellular carcinoma (HCC) ^[1]^. Alcoholic fatty liver (AFL) has been considered as critical pathophysiological processing to develop HS and HC^[2]^. Immoderate alcohol drinking pattern are exposed to alcoholic fatty liver disease (AFLD), hepatic fibrosis (HF), and hepatitis in the same patient groups consecutively ^[1,3]^. HS is an initial indication to acute alcoholism, which is a disorder occurred by excessive accumulation of fat in hepatocytes ^[4]^. HS can progress steatohepatitis, which triggers extreme inflammation in liver ^[5]^. HS can develop into HF, which activates hepatic stellate cells (HSCs) via the gradual thickening of extracellular matrix (ECM) ^[6]^. The abnormal alteration caused by ECM construction and the activation of HSCs gives birth to HF. Commonly, HF is an initial stage of liver injury, the progression of which can spawn HC ^[7]^. Typically, HC is the most critical risk factor to be developed into HCC ^[8]^. Hence, the etiological development of ALD leads to liver failure as irreparable stage. As a rule, liver injury caused by alcohol may be restored if abstention is sustained persistently, evidently, alcohol withdrawal is an optimal therapeutic strategy against ALD ^[9]^. At present, as another substitute, metadoxine has been administered to lower alcohol amount in the blood, however, causes some negative side effects: epigastric burning, nausea, and vomiting ^[10,11]^.

Because of these downsides, medicinal plants are significant resources to resolve unexpected side effects against human disease ^[12]^. Based on this, *Gleditsia sinensis Lam.* (GSL) has been valued as therapeutic herb used for anti-hyperlipidemic activity ^[13]^. Another report demonstrated that GSL dampens lipid accumulation and inhibits the adipogenic transcription factors including cell cycle arrest ^[14]^. In spite of these effects, the key molecules and exact mechanism of GSL is yet to be documented. Accordingly, the aim of this study is to obtain the therapeutic gain from GSL in the treatment of ALD.

Most recently, gut microbiota (GM) treatment come into spotlight as another therapeutic resource to enable to provide beneficial efficacy for fatty liver disease (FLD) patients ^[15]^. In prior studies, various metabolites produced by GM might be mediators to alleviate FLD, for instance, by modulating the pharmacological mode(s) associated with immune response ^[16,17]^. The indoles and its derivatives produced by GM modulate immune homeostasis and metabolic disorders related to insulin resistance ^[18]^. Also, the polyphenols produced by GM have favorable effects on the host-GM interactions: anti-oxidation and anti-inflammation ^[19]^. Collectively, we integrated the significant exogenous species (or GSL) and endogenous species (or GM) to unravel their interrelationships. This newly emerged platform can be allowed to discover new agents against ALD. In 2007, Hopkins first defined as network pharmacology (NP), an integrated-based drug discovery that combines bioinformatics, cheminformatics, and system biology ^[20]^. The emergence of NP has been changed the research concept from “single target, single molecule, single mechanism” to “multiple targets, multiple molecules, multiple mechanisms” by exposing collective components target(s)-molecule(s) and target(s)-mechanism(s) networks to help elucidate the reasonableness and adaptation of medication ^[21]^.

In this study, we updated the NP platform with microbial informatics that included targets, metabolites from GM, which was devised to investigate the relationships between exogenous species (GSL) and endogenous species (GM). Correspondingly, we constructed GSL or GM-Signaling pathways – Targets – Metabolites (GGSTM) networks to decrypt its relationships. The study processing was briefly represented as below. Firstly, methanolic extraction (ME) GSL was analyzed by Gas chromatography mass spectrum (GC-MS) to obtain bioactive compounds, which were filtered by Lipinski’s rule to select the drug-like molecules. The verified molecules were retrieved its targets through public databases, then, the intersecting targets between the two databases were gathered. The ALD-responded targets were acquired by human-based public bioinformatics and the common targets were selected to achieve the exactness and robustness. Secondly, the gathered overlapping targets were considered as crucial targets, suggesting that protein -protein interaction (PPI) networks and a bubble chart were required to identify the substantial elements. Thirdly, microbial informatics to browse the key factors related to GM was employed to construct GGSTM networks. Then, the final represented targets were subjected to molecular docking test (MDT) and density functional theory (DFT) to affirm the affinity and reactivity of corresponding molecules, the results of which could be determined as the most promising effectors in the treatment of ALD. The detailed information was exhibited in **Figure 1**.

**Figure 1.**
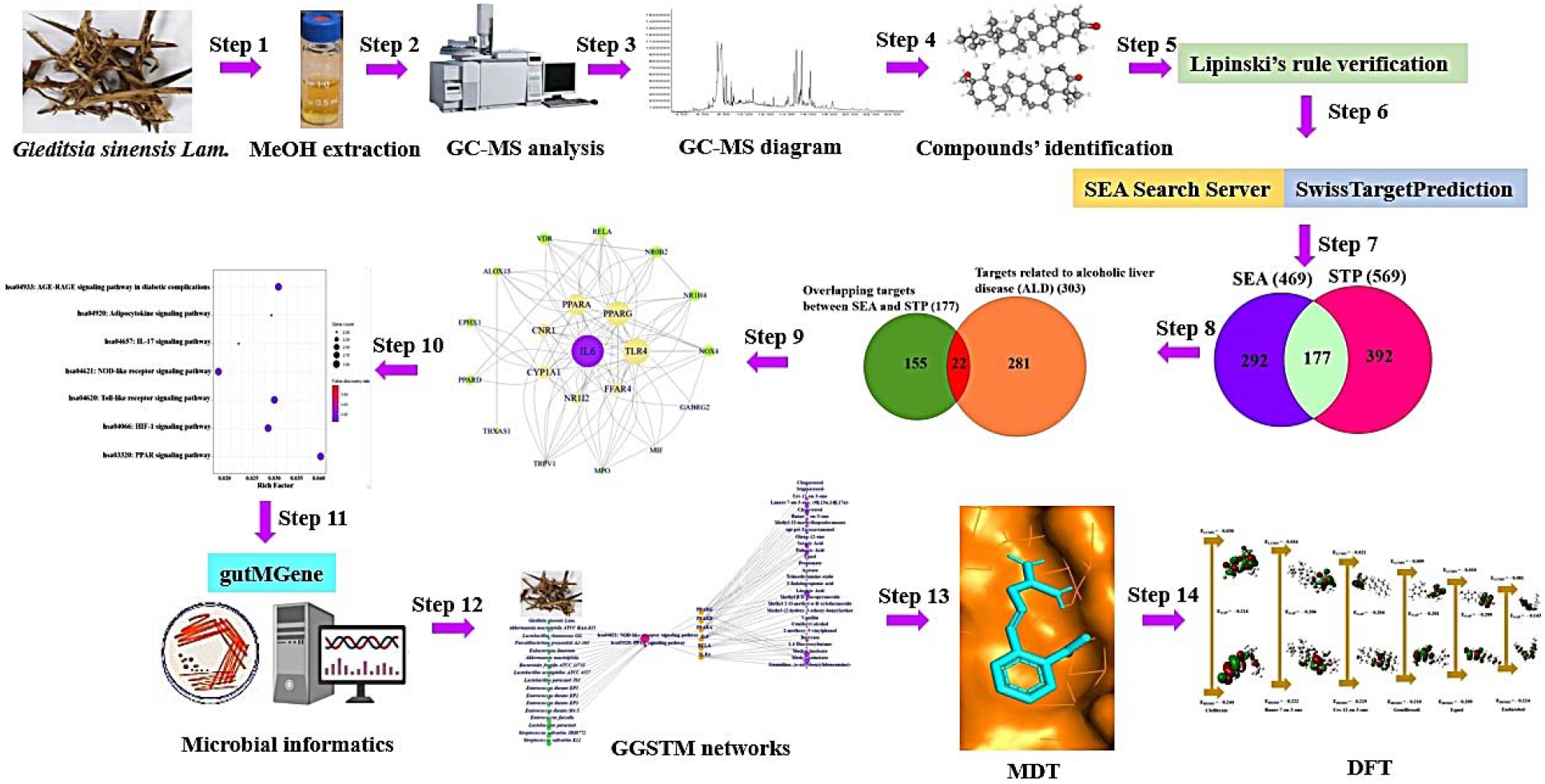
The schematic process of this study.

## Results

### The identification of chemical constituents from GSL

A total of 47 chemical constituents were detected through Gas Chromatography Mass Spectrum (GC-MS) (**Figure 2B**), profiling up the distinct information of each compound, including chemical taxonomy (**Table EV1**). The number of 47 chemical constituents were confirmed by Lipinski’s rule (Molecular Weight ≤ 500 g/ mol; Moriguchi octanol–water partition coefficient ≤ 4.15; Number of Nitrogen or Oxygen ≤ 10; Number of NH or OH ≤ 5). The Topological Polar Surface Area (TPSA) value was considered as propensity to be determined as cell permeability, the cut-off of which is less than 140 Å^2^ ^[28]^. As mentioned above, drug-like molecules (DLMs) were accepted by its filtering methodology, and the detailed contents were listed up in **Table 1**.

**Figure 2.**
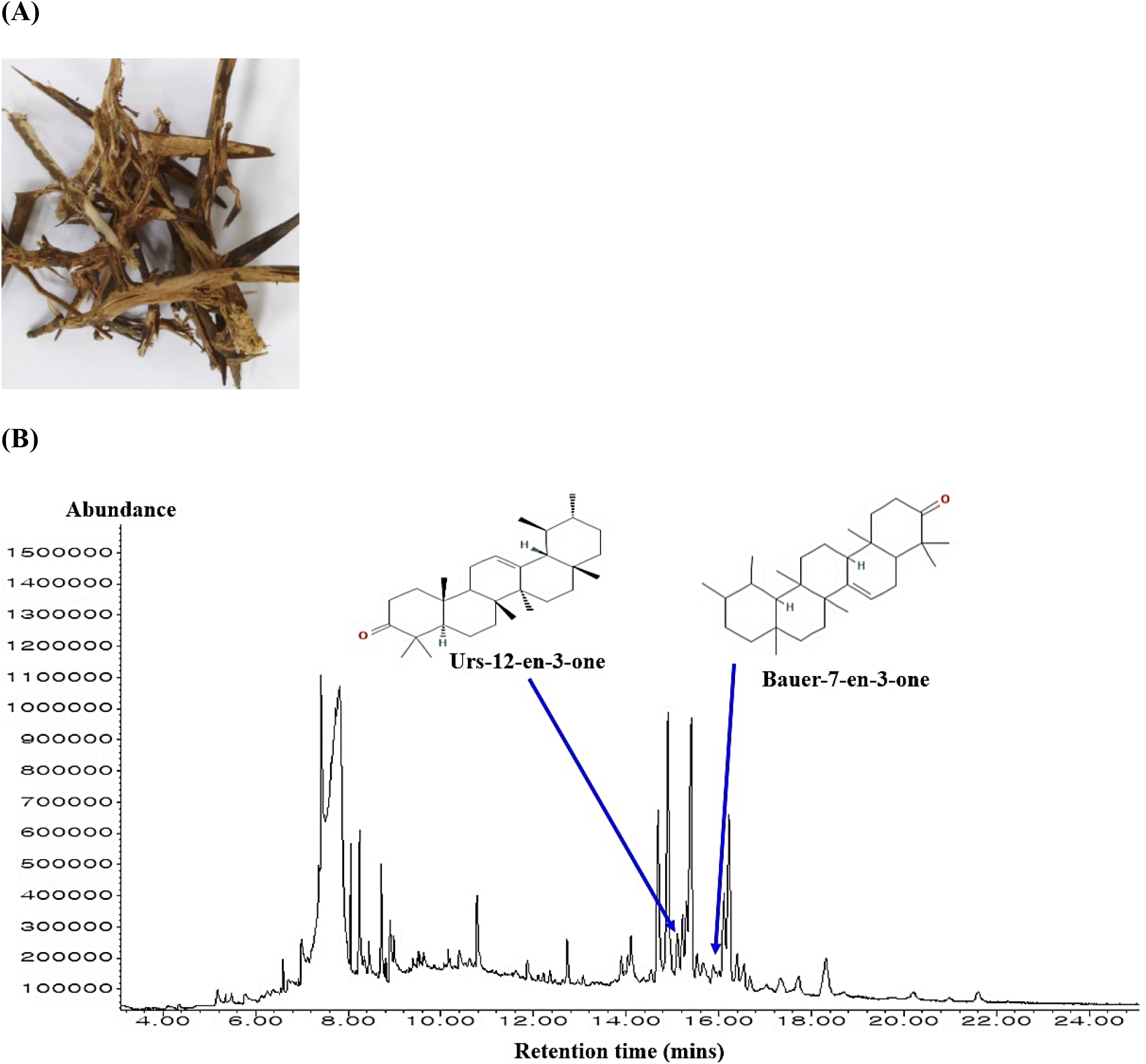
(A) The picture of GSL. (B) GC-MS chromatography of methanolic extract of GSL and key molecules.

**Table 1.** The physicochemical properties of molecules from GSL.

| No. | Compound name | PubChem CID | Lipinski Rules |  |  |  | Lipinski's violations | Bioavailability Score | TPSA |
| --- | --- | --- | --- | --- | --- | --- | --- | --- | --- |
|  |  |  | MW | HBA | HBD | MLogP |  |  |  |
|  |  |  | < 500 | < 10 | ≤ 5 | ≤ 4.15 | ≤ 1 | > 0.1 | < 140 Å² |
| 1 | 2-methoxy-5-vinylphenol | 10441857 | 150.17 | 2 | 1 | 1.71 | 0 | 0.55 | 29.46 |
| 2 | 1,4-Diacetoxybutane | 69410 | 174.19 | 4 | 0 | 0.93 | 0 | 0.55 | 52.60 |
| 3 | Carbamic acid, dimethyl-, ethyl ester | 12709 | 117.15 | 2 | 0 | 0.42 | 0 | 0.55 | 29.54 |
| 4 | Vanillin | 1183 | 152.15 | 3 | 1 | 0.51 | 0 | 0.55 | 46.53 |
| 5 | Kodaflex txib | 23284 | 286.41 | 4 | 0 | 3.17 | 0 | 0.55 | 52.60 |
| 6 | Methyl-(2-hydroxy-3-ethoxy-benzyl)ether | 586449 | 182.22 | 3 | 1 | 1.22 | 0 | 0.55 | 38.69 |
| 7 | Pyridine, 4-ethyl- | 10822 | 107.15 | 1 | 0 | 1.15 | 0 | 0.55 | 12.89 |
| 8 | Ethane, isothiocyanato- | 10966 | 87.14 | 1 | 0 | 1.62 | 0 | 0.55 | 44.45 |
| 9 | Coniferyl alcohol | 1549095 | 180.20 | 3 | 2 | 1.13 | 0 | 0.55 | 49.69 |
| 10 | Methyl beta-D-glucopyranoside | 445238 | 194.18 | 6 | 4 | -2.40 | 0 | 0.55 | 99.38 |
| 11 | Methyl 2-O-methyl-alpha-D-xylofuranoside | 12775307 | 178.18 | 5 | 2 | -1.59 | 0 | 0.55 | 68.15 |
| 12 | Methyl palmitate | 8181 | 270.45 | 2 | 0 | 4.44 | 1 | 0.55 | 26.30 |
| 13 | Palmitic Acid | 985 | 256.42 | 2 | 1 | 4.19 | 1 | 0.85 | 37.30 |
| 14 | Methyl linoleate | 5284421 | 294.47 | 2 | 0 | 4.70 | 1 | 0.55 | 26.30 |
| 15 | Methyl 15-methylheptadecanoate | 554152 | 298.50 | 2 | 0 | 4.91 | 1 | 0.55 | 26.30 |
| 16 | Linoleic Acid | 5280450 | 280.45 | 2 | 1 | 4.47 | 1 | 0.85 | 37.30 |
| 17 | Stearic Acid | 5281 | 284.48 | 2 | 1 | 4.67 | 1 | 0.85 | 37.30 |
| 18 | 7-(1,3-Dimethylbuta-1,3-dienyl)-1,6,6-trimethyl-3,8-dioxatricyclo[5.1.0.0(2,4)]octane | 5363696 | 234.33 | 2 | 0 | 2.63 | 0 | 0.55 | 25.06 |
| 19 | Vulgarol A | 91746444 | 308.50 | 2 | 2 | 3.93 | 0 | 0.55 | 40.46 |
| 20 | Isohaloethane | 9628 | 197.38 | 3 | 0 | 2.60 | 0 | 0.55 | 0.00 |
| 21 | Cannithrene 1 | 53438738 | 242.27 | 3 | 2 | 2.20 | 0 | 0.55 | 49.69 |
| 22 | Nitroscanate | 68547 | 272.28 | 4 | 0 | 3.30 | 0 | 0.55 | 99.50 |
| 23 | 3-Hydroxycarbofuran phenol | 86626 | 180.20 | 3 | 2 | 0.83 | 0 | 0.55 | 49.69 |
| 24 | Pregnan-20-one # | 1715430 | 302.49 | 1 | 0 | 5.08 | 1 | 0.55 | 17.07 |
| 25 | Longiborneol | 6432447 | 222.37 | 1 | 1 | 3.81 | 0 | 0.55 | 20.23 |
| 26 | Cholesterol | 5997 | 386.65 | 1 | 1 | 6.34 | 1 | 0.55 | 20.23 |
| 27 | Cholesterol methyl ether | 262105 | 40.68 | 1 | 0 | 6.54 | 1 | 0.55 | 9.23 |
| 28 | DMD | 7210 | 282.34 | 4 | 2 | 1.81 | 0 | 0.55 | 65.18 |
| 29 | Ursa-9(11),12-dien-3-one | 634574 | 422.69 | 1 | 0 | 6.72 | 1 | 0.55 | 17.07 |
| 30 | Stigmasterol | 5280794 | 412.69 | 1 | 1 | 6.62 | 1 | 0.55 | 20.23 |
| 31 | 3-Epicycloecalenol | 543796 | 426.72 | 1 | 1 | 6.92 | 1 | 0.55 | 20.23 |
| 32 | Clionasterol | 457801 | 414.71 | 1 | 1 | 6.73 | 1 | 0.55 | 20.23 |
| 33 | Olean-12-ene | 11058690 | 410.72 | 0 | 0 | 8.91 | 1 | 0.55 | 0.00 |
| 34 | Cycloecalenol | 101690 | 426.72 | 1 | 1 | 6.92 | 1 | 0.55 | 20.23 |
| 35 | Oprea1_549827 | 627194 | 300.40 | 1 | 0 | 3.88 | 0 | 0.55 | 17.30 |
| 36 | Bauer-7-en-3-one | 618607 | 424.70 | 1 | 0 | 6.82 | 1 | 0.55 | 17.07 |
| 37 | Urs-12-en-3-one | 124018 | 424.70 | 1 | 0 | 6.82 | 1 | 0.55 | 17.07 |
| 38 | Cycloartenol acetate | 518616 | 468.75 | 2 | 0 | 7.08 | 1 | 0.55 | 26.30 |
| 39 | 4,6,6-trimethyl-2-[(1E)-3-methylbuta-1,3-dienyl]-3-oxatricyclo[5.1.0.0(2,4)]octane | 5369926 | 218.33 | 1 | 0 | 3.56 | 0 | 0.55 | 12.53 |
| 40 | 2-Methyl-6-(5-methyl-2-thiazolin-2-ylamino)pyridine | 610051 | 207.30 | 2 | 1 | 1.69 | 0 | 0.55 | 62.58 |
| 41 | Stigmast-4-en-3-one | 5484202 | 412.69 | 1 | 0 | 6.62 | 1 | 0.55 | 17.07 |
| 42 | epi-psi-Taraxastanonol | 101290252 | 442.72 | 2 | 1 | 6.00 | 1 | 0.55 | 37.30 |
| 43 | STK691054 | 610051 | 207.30 | 2 | 1 | 1.69 | 0 | 0.55 | 62.58 |
| 44 | Ergosta-14,22-dien-3-ol, acetate, (3.beta.,5.alpha.,22E)- | 319424149 | 440.70 | 2 | 0 | 6.61 | 1 | 0.55 | 26.30 |
| 45 | PTH-DL-proline | 100798 | 232.30 | 1 | 0 | 1.77 | 0 | 0.55 | 55.64 |
| 46 | Lanost-7-en-3-one, (9beta,13alpha,14beta,17alpha)- | 91691424 | 426.72 | 1 | 0 | 6.82 | 1 | 0.55 | 17.07 |
| 47 | Guanidine, (o-nitrobenzylideneamino)- | 9562955 | 207.19 | 4 | 2 | 0.10 | 0 | 0.55 | 122.58 |

### Overlapping targets between SEA and STP databases

The targets related to DLMs were gathered by SEA (469) and STP (569) databases, suggesting that the identified targets are significant elements from GSL (**Table EV2**). The number of 177 overlapping targets was selected to achieve the rigorousness from the two databases (**Table EV2**) (**Figure 3A**).

**Figure 3.**
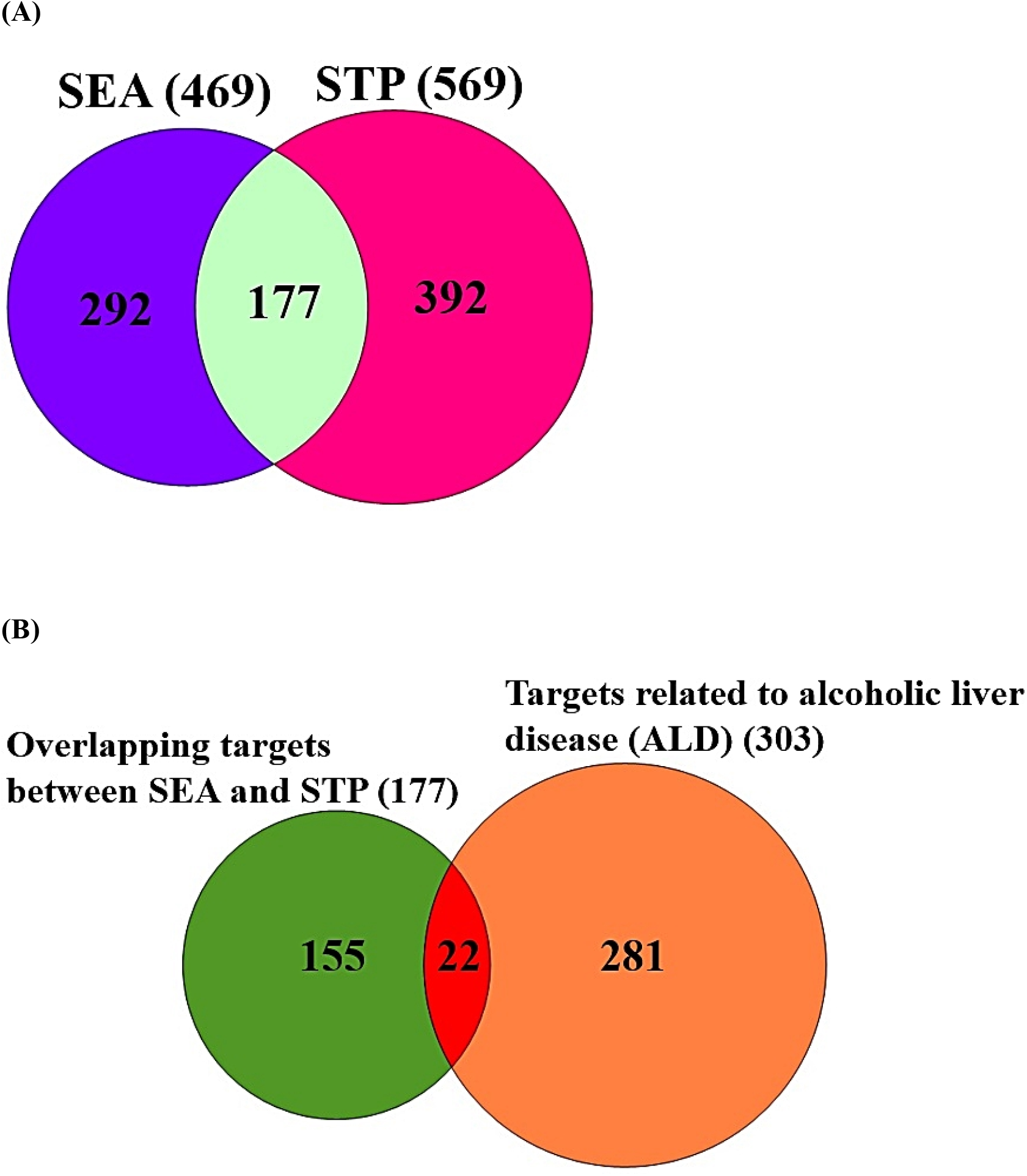
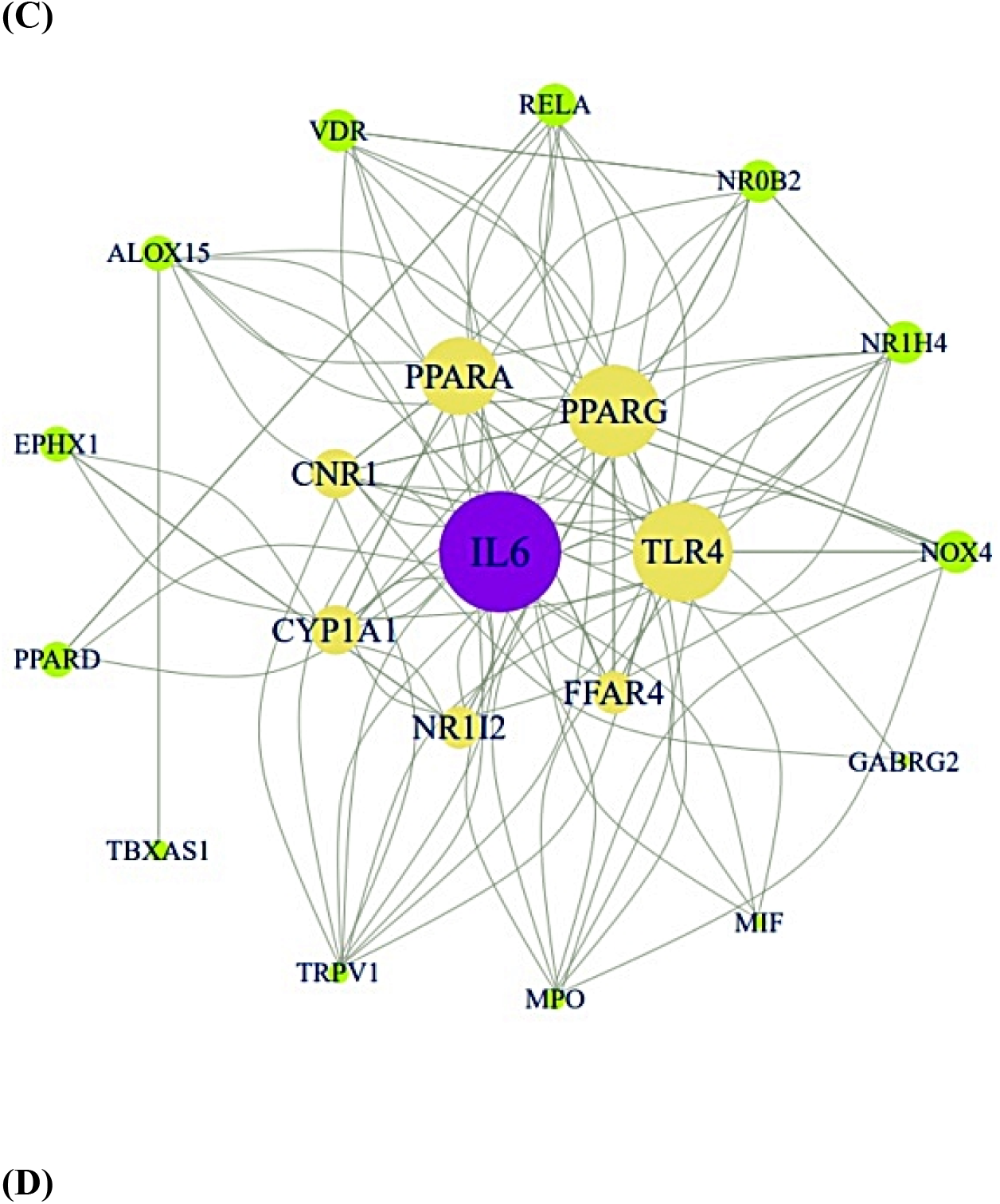
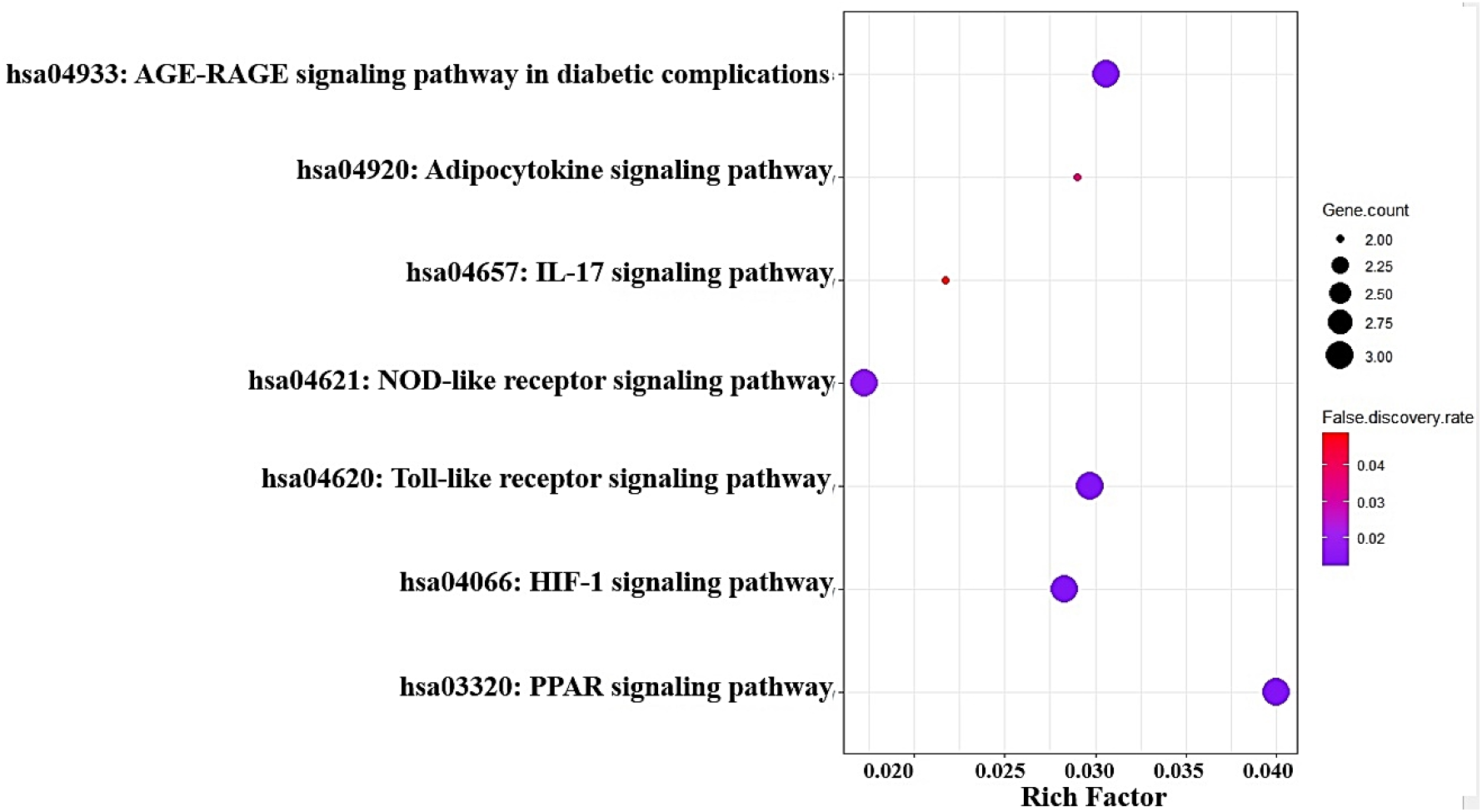
(A) The number of overlapping 177 targets between SEA (469) and STP (569). (B) The number of 22 final targets between 177 targets and ALD-related targets (303). (C) The PPI networks (21 nodes, 118 edges). (D) A bubble plot of the 7 signaling pathways to be associated with ALD.

### Identification of ALD-related targets and key targets

Two public databases (DisGeNET, and OMIM) yielded a total of 303 targets related to ALD (**Table EV2**). An input term to gather the targets was “alcoholic liver disease” in the two databases. The Venn diagram plotter showed the 22 overlapping targets (**Table EV2**) between ALD-related targets (303) and 177 GSL–related targets (**Figure 3B**). The final 22 targets were considered as key targets in treating ALD.

### PPI networks and a bubble plot

The significant 22 targets were added to STRING database to extract source of the PPI networks to elucidate potential pathways of GSL anti-ALD action. The STRING network analysis demonstrated that all targets in the network related to each other via 21 nodes, 118 edges (**Figure 3C**). In 22 targets, GSTK1 targets had no correlation with other targets. R Package was employed to obtain the DV as the significant level in the networks. As a result, IL6 had the highest DV (32), which was followed by TLR4 (26), PPARG (24), and PPARA (20) (**Table EV3**). Accordingly, IL6 is a control tower to modulate the neighbor targets associated with ALD targets. Based on STRING network, a bubble plot was depicted to represent 7 signaling pathways, thereby, NOD-like receptor signaling pathway with the lowest rich factor might function as antagonistic mechanism, and PPAR signaling pathway with the highest rich factor might work as agonistic mode in alleviating ALD (**Figure 3D**) (**Table EV4**). Thus, the two signaling pathways might be amenable to regulate the pathophysiology of ALD.

### The connectivity to GSL and GM via GGSTM networks

Based on GSL, 16 GM, 2 signaling pathways, 6 targets, and 29 metabolites (including 9 GM’s metabolites), GGSTM networks were constructed by the amount of connectivity on each element. Correspondingly, the GGSTM networks ascertain that GSL molecules are mediators to inhibit NOD-like receptor signaling pathway, GM’s metabolites are mediators to activate PPAR signaling pathway, which can function as dual agents in treating ALD. The GGSTM networks comprised 54 nodes (1 GSL, 16 GM, 2 signaling pathways, 6 targets, and 29 metabolites) and 74 edges. More extensively, GSL, *Akkermansia muciniphila ATCC BAA-835*, *Lactobacillus rhamnosus GG*, *Faecalibacterium prausnitzii A2–165*, *Eubacterium limosum, Akkermansia muciniphila*, *Bacteroides fragilis ATCC 23745*, *Lactobacillus acidophilus ATCC 4357*, *Eubacterium limosum*, *Lactobacillus paracasei JS1*, *Enterococcus durans EP1*, *Enterococcus durans EP2*, *Enterococcus durans EP3*, and *Enterococcus durans M4-5* are linked to NOD-like signaling pathways on IL6, RELA, and TLR4 with 17 molecules (or metabolites). Then, *Enterococcus faecalis*, *Lactobacillus paracasei*, *Streptococcus salivarius JIM8772*, and *Streptococcus salivarius K12* are related to PPAR signaling pathway on PPARA, PPARD, and PPARG with 15 molecules (or metabolites) (**Figure 4**). In a total of 29 metabolites, Linoleic acid, Methyl palmitate, and Methyl linoleate are involved in the two key signaling pathways responded to ALD. Thus, GSL molecules and GM’s metabolites show intimate relationships to modulate in treating ALD.

**Figure 4.**
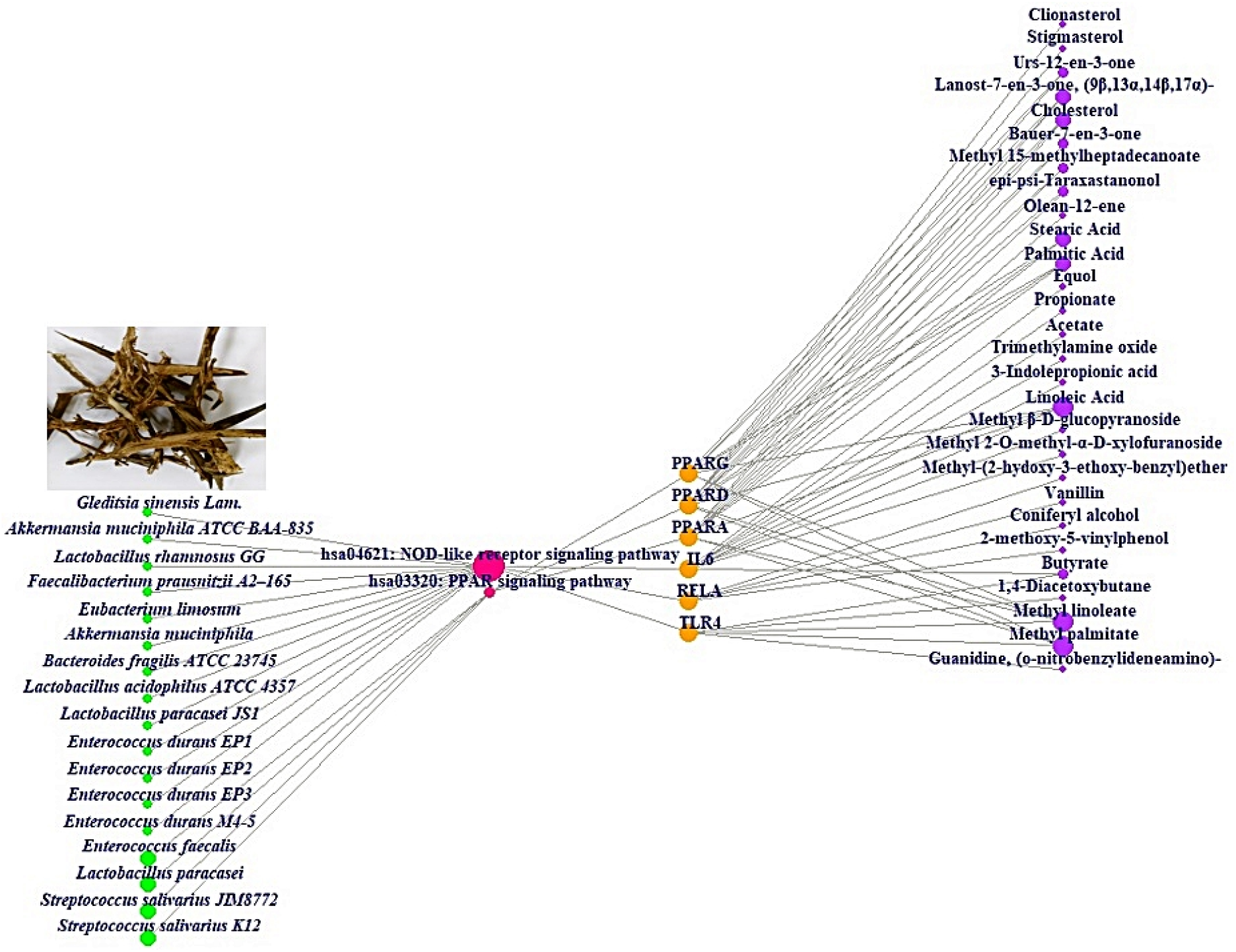
GSL or GM – Signaling pathways – Targets – Metabolites (GGSTM) networks (54 nodes and 74 edges). Green color circle: GM; Red color circle: signaling pathway; Orange color circle: target; Purple color circle: metabolite.

### Molecular docking test (MDT) assessment

In the NOD-like receptor signaling pathway, the MDT revealed that Equol – IL6 (PDB ID: 3SP6) conformer formed the most stable complex with -7.4 kcal/mol (**Figure 5A**). Noticeably, Equol was attributed to *Lactobacillus paracasei JS1* and any other targets (TLR4, and RELA) had not valid docking score (≤ -6.0kcal/mol) ^[29]^. Furthermore, LMT-28 ^[30]^ as a representative IL6 inhibitor was compared with Equol to evaluate what the molecule with better binding affinity is. As a result, Equol was more optimal molecule than LMT-28 (– 6.2 kcal/mol) on IL6.

**Figure 5.**
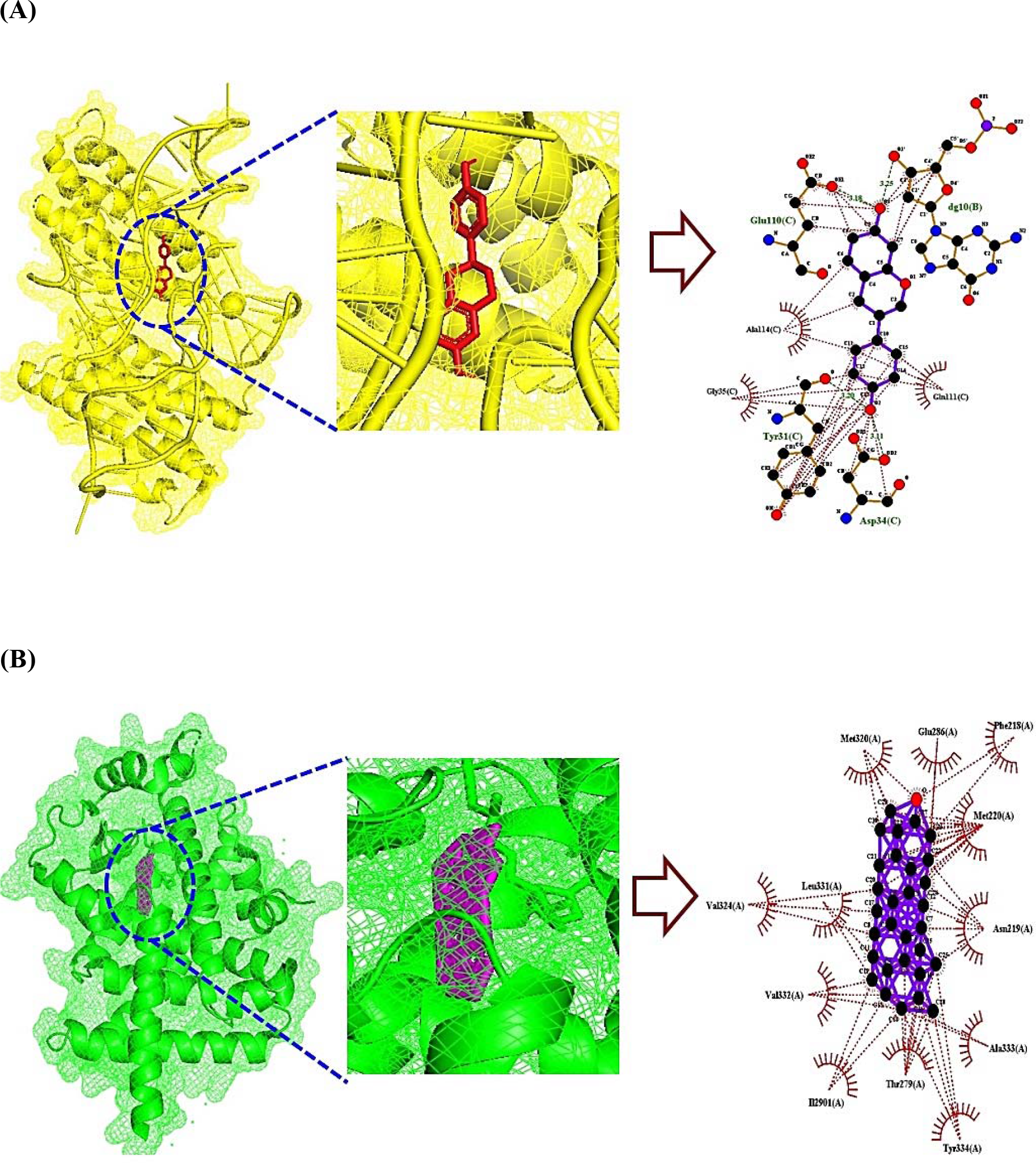
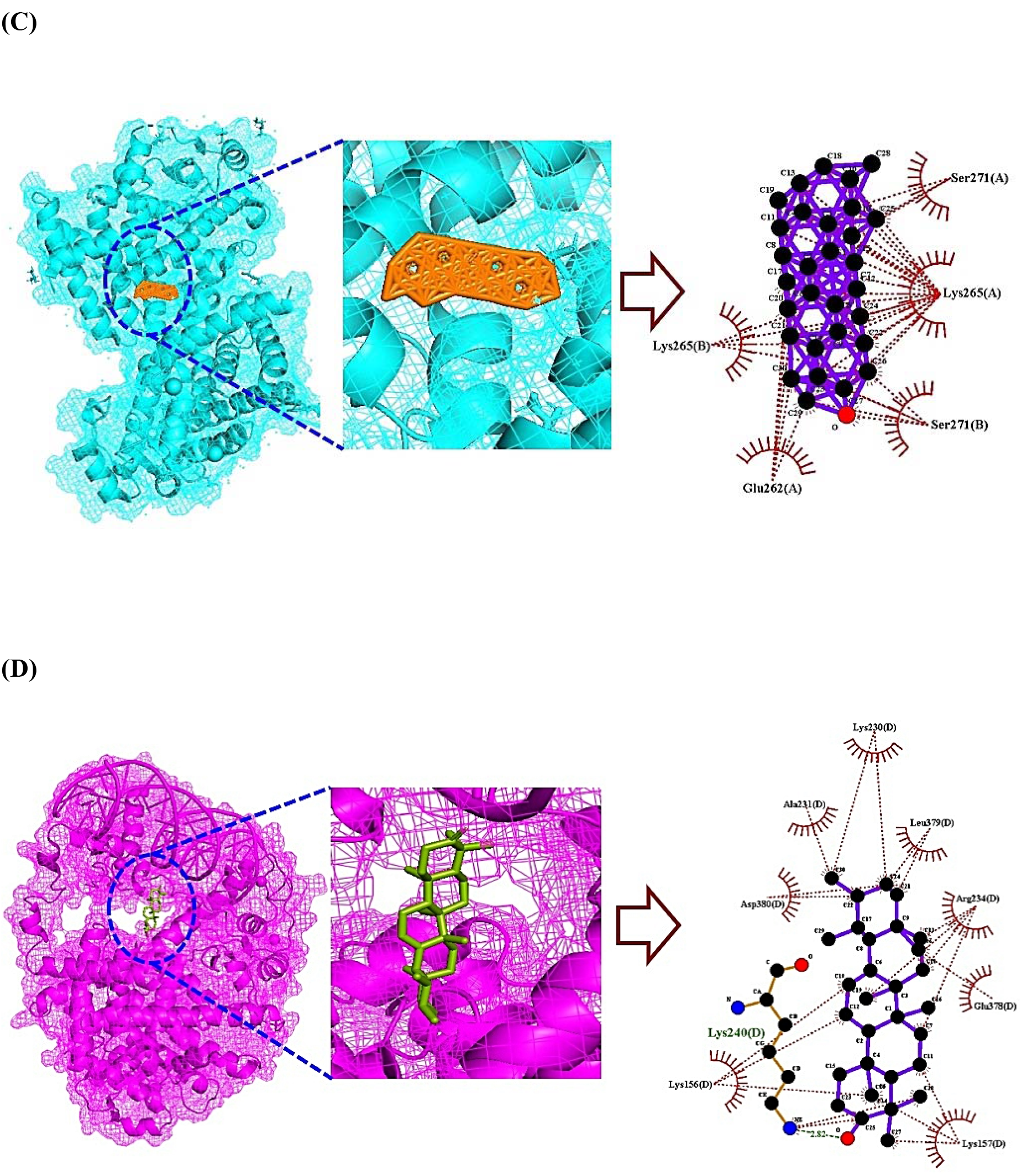
The 2-dimension (2D), 3-dimension (3D) representation of Molecular docking test (MDT) between key targets and molecules. (A) IL6 (PDB ID: 4NI9) – Equol; Binding affinity: – 7.4 kcal/mol (B) PPARA (PDB ID: 3SP6) – Bauer–7–en–3–one; Binding affinity: – 16.3 kcal/mol. (C) PPARD (PDB ID: 5U3Q) – Bauer–7–en–3–one; Binding affinity: – 14.1 kcal/mol. (D) PPARG (PDB ID: 3E00) – Urs-12-en-3-one; Binding affinity: – 8.7 kcal/mol.

In the PPAR signaling pathway, the MDT demonstrated that Bauer-7-en-3-one – PPARA (**Figure 5B**), Bauer-7-en-3-one – PPARD (**Figure 5C**), and Urs-12-en-3-one – PPARG (**Figure 5D**) conformers formed the most stable complex with -16.3 kcal/mol, -14.1 kcal/mol, and -8.7 kcal/mol, respectively.

Specifically, the uppermost two molecules (Bauer-7-en-3-one, and Urs-12-en-3-one) were originated from GSL. In comparison with 5 positive controls (Clofibrate ^[31]^ on PPARA, Endurobol ^[32]^ on PPARD, and Pioglitazone ^[33]^, Rosiglitazone ^[34]^ on PPARG), Bauer-7-en-3-one, and Urs-12-en-3-one from GSL were greater binding affinity than the known standard drugs. The detailed information was profiled in **Table EV5**.

### DFT comparison between key molecules and standard drugs

The DFT of key molecules and standard drugs were adopted to investigate their physicochemical reactivity on certain targets. Commonly, how a molecule functions as a donator or as acceptor on its valence electrons to another molecule can be assessed by the HOMO and LUMO values. Herein, the three key molecules (Bauer-7-en-3-one, Urs-12-en-3-one, and Equol) showed substantial HOMO energy levels (–0.222 kcal/mol, –0.225 kcal/mol, and –0.209 kcal/mol) compared to Clofibrate, suggesting that increasing conjugation boosted the HOMO energy with electron donator groups, however, decreased the LUMO energy. Namely, the three key molecules were good electron donators by comparison with the standard drugs. In a way, the stability of a molecule can be confirmed by Energy gap (E_GAP_) (HOMO – LUMO) and lower E_GAP_ is considered as a soft effector (**Figure 6).** Collectively, the three key molecules had certain valid parameters, including LUMO, HOMO, E_GAP_, Hardness (ɳ), Softness (S), and electronegativity (χ) (**Table 2)**.

**Figure 6.**
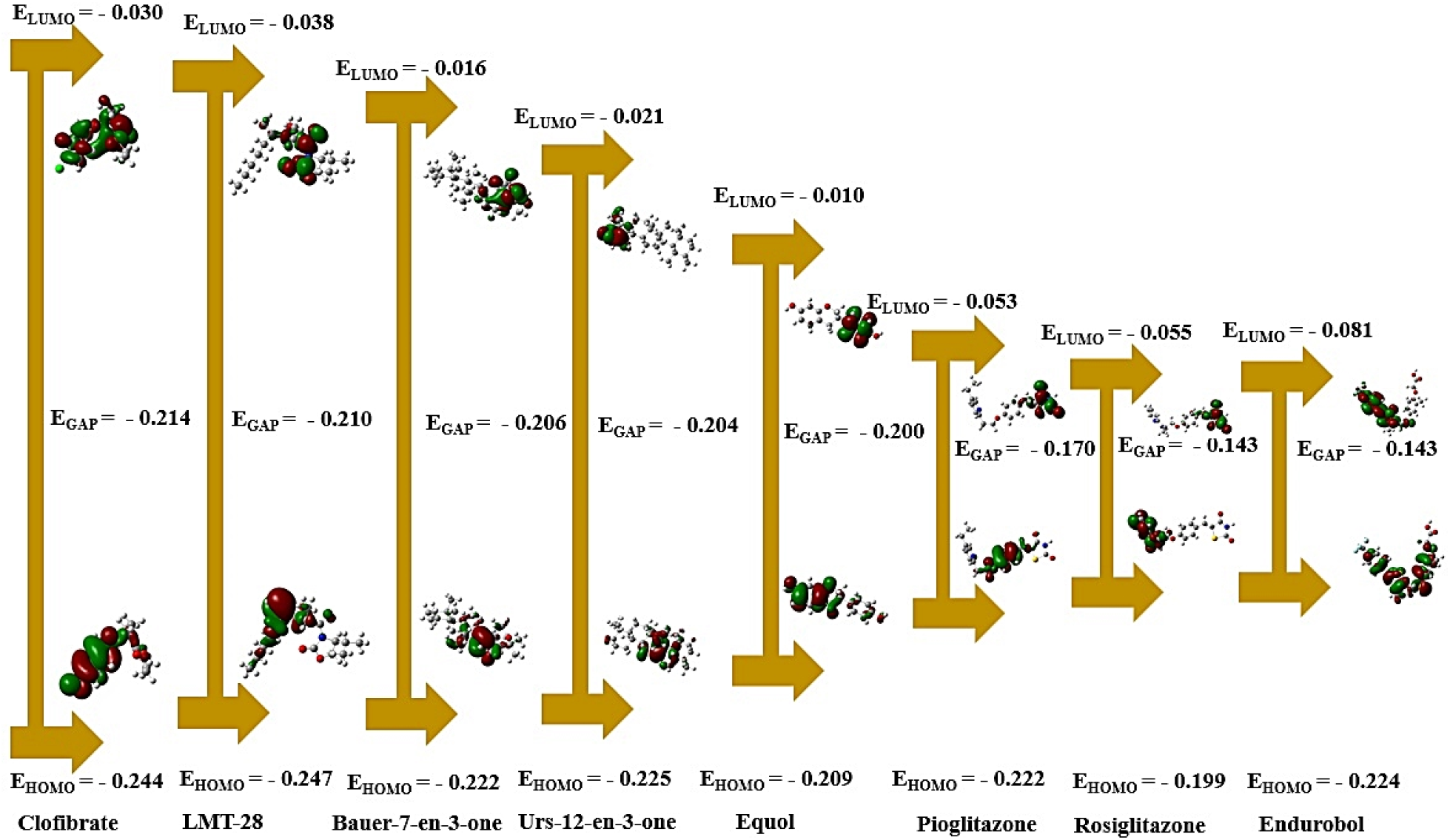
The LUMO – HOMO values and the E_GAP_ of molecular orbital for key molecules from GSL and standard drugs (Clofibrate, LMT-28, Pioglitazone, Rosiglitazone, and Endurobol). Green color orbitals: Negative phase; Red color orbitals: Positive phase.

**Table 2.** The calculated parameters on frontiers molecular orbital (FMO) theory.

| No. | Molecules | LUMO | HOMO | $E_{\text{GAP}}$ (eV) | $\eta$ (eV) | S (eV) | $\chi$ (eV) |
| --- | --- | --- | --- | --- | --- | --- | --- |
| 1 | Clofibrate | -0.030 | -0.244 | -0.214 | 0.107 | 9.352 | -0.107 |
| 2 | LMT-28 | -0.038 | -0.247 | -0.210 | 0.105 | 9.542 | -0.105 |
| 3 | Bauer-7-en-3-one | -0.016 | -0.222 | -0.206 | 0.103 | 9.690 | -0.103 |
| 4 | Urs-12-en-3-one | -0.021 | -0.225 | -0.204 | 0.102 | 9.786 | -0.102 |
| 5 | Equol | -0.010 | -0.209 | -0.200 | 0.100 | 10.025 | -0.100 |
| 6 | Pioglitazone | -0.053 | -0.222 | -0.170 | 0.085 | 11.783 | -0.085 |
| 7 | Rosiglitazone | -0.055 | -0.199 | -0.143 | 0.072 | 13.951 | -0.072 |
| 8 | Endurobol | -0.081 | -0.224 | -0.143 | 0.072 | 13.974 | -0.072 |

## Discussion

The GSL is known as a significant medicinal herb to relieve ALD; however, its key bioactives, targets, and mechanism(s) are yet to be clarified completely. We employed GC-MS analysis to identify chemical compounds with druglike properties, indicating that the detected compounds can be permeable to cell membranes to reach certain targets ^[35]^. In parallel, these chemical compounds were considered as effectors against ALD. In another scenario, GM is considerable element to alleviate ALD, and its metabolites contribute to exert the therapeutic value ^[36]^. The aim of this study was to establish the favorable events by incorporating the components of GSL and GM in treating of ALD. This methodology has been introduced firstly to reveal the relationships between GSL and GM. To complete this project, we integrated the concept of bioinformatics, cheminformatics, pharmacology, microbiology, natural products chemistry, and quantum chemistry to enhance the scientific evidence in entangled microbiome atlas. Furthermore, we employed the DFT theory to confirm the potentiality of drug-like property, which was manifested the stability and reactivity of key molecules compared with standard drugs (LMT-28, Clofibrate, Endurobol, Pioglitazone, and Rosiglitazone). All theoretical parameters to characterize their intrinsic property were obtained at minimum energy optimization, which are crucial factors to determine the greater chemical stability and binding affinity, more importantly, Hardness (ɳ), and Softness (S) ^[37–39]^.

Ultimately, molecular theory (MT) was employed to identify the medicinal value of key molecules, describing that the corresponding parameters are important assessors against the standard drugs. In this study, we established the combinatorial effects against ALD via NOD-like receptor signaling pathway, and PPAR signaling pathway. The uppermost target in PPI networks was IL6 controlled by Equol as a metabolite from *Lactobacillus paracasei JS1,* which was linked to NOD-like receptor signaling pathway. Herein, NOD-like receptor signaling pathway with lowest rich factor represented by bubble chart was attributed to antagonistic mode. In parallel, IL6 as a key target, *Lactobacillus paracasei JS1* as a favorable GM, and Equol as a key metabolite were all significant components on NOD-like receptor signaling pathway in treating ALD. A study to support this result has been reported that NOD-like receptor activated in the liver injury, which was caused hepatocellular cytotoxicity ^[40]^. It elicits that inhibition of NOD-like receptor signaling pathway can make favorable effects on ALD via dampening inflammasome.

Knowingly, agonists of PPAR signaling pathway play central role in alleviating the inflammatory responses in hepatocytes ^[41]^. Noticeably, Bauer-7-en-3-one and Urs-12-en-3-one from GSL were potent effectors in MDT although the metabolites from *Enterococcus faecalis*, *Lactobacillus paracasei*, *Streptococcus salivarius JIM8772*, and *Streptococcus salivarius* are involved in PPAR signaling pathway activation. The two molecules are categorized into triterpenoid species, suggesting that they possess antioxidant, metabolic regulator, immunomodulator, anti-inflammatory, anti-cancer efficacy including hepatoprotective agents ^[42]^. It infers that the two triterpenoids might be potential agents to thwart hepatic injury from ALD.

MDT revealed that Equol, Bauer-7-en-3-one and Urs-12-en-3-one had the strongest binding affinity on IL6, PPARA, PPARD, and PPARG, compared with standard drugs (LMT-28, Clofibrate, Endurobol, Pioglitazone, and Rosiglitazone). It means that the three molecules with the lowest binding energy are more stable binding partners on targets ^[43]^.

Furthermore, frontier molecular orbital (FMO) analysis was performed to confirm the molecule’s physicochemical reactivity. The intrinsic stability and reactivity depends on E_GAP_ via LUMO and HOMO ^[44]^. The great kinetic stability and physicochemical reactivity may be characterized by E_GAP_, indicating that E_GAP_ is directed to decrypt the hardness and softness ^[45]^. The higher softness level boosts its reactivity between binding partner(s) and our pioneered three molecules had valid parameters to develop as new agents. Noticeably, Bauer-7-en-3-one and Urs-12-en-3-one demonstrated a higher value of softness than Clofibrate, suggesting that the two triterpenoids from GSL might be alternatives to activate PPAR signaling pathway.

Evidently, Bauer-7-en-3-one on PPARA had more valid value than PPARD, and PPARG in aspects of MDA and FMO analysis. This is the reason why Bauer-7-en-3-one shows better binding affinity and reactivity than a standard drug (Clofibrate) as PPARA agonist. Although Bauer-7-en-3-one had more stable binding affinity than PPARD, and PPARG, in comparison with standard drugs (Endurobol, Pioglitazone, and Rosiglitazone), which was less reactivity level in FMO analysis. To be pinpointed, Bauer-7-en-3-one is more optimal for a new PPARA agonist in viewpoint of pan-PPAR agonism. All things considered, these findings suggest that Equol, Bauer-7-en-3-one, and Urs-12-en-3-one would be multiple potential effectors to relieve ALD.

### The opportunities and obstacles of this study

The primary aim of this study was to integrate key therapeutic-related components of GSL and GM in treating ALD. The scheme has been centered on uncovering the relationships between GSL and GM, which was arranged as a platform mode to enter new information without any hurdle. To complete this project, we devised a novel scenario to pursue the exactness and rigorousness concerning biological output. In addition, we adopted the concept of orbital theory to realize drug-like value of key molecules, compared with standard drugs. In this study, we combined some academic fields (bioinformatics, cheminformatics, pharmacology, microbiology, natural products chemistry, analytical chemistry, and quantum chemistry) to discover therapeutic results against ALD. The setup might be a hallmark to unravel complicated interrelationships between GSL and GM, extensively, the application of which can be extended to exogenous species (food) – endogenous species (GM) – host system.

This study shows that end-products achieved by the platform are important components to establish the combinatorial effects via incorporation of GSL and GM for anti-ALD. On holistic viewpoint, the setup is a debut version to analyze ALD in microbiome research and is a sound methodology to discover new findings. Nonetheless, more *in vivo* data and its pharmacokinetic studies are required to obtain better clarification on the complex pattern against ALD.

## Conclusion

To sum up, this study represents that IL6-Equol conformer from *Lactobacillus paracasei JS1* as antagonism on NOD-like receptor signaling pathway, PPARA–Bauer-7-en-3-one; PPARD– Bauer-7-en-3-one; and PPARG-Urs-12-en-3-one conformers from GSL as agonism on PPAR signaling pathway in ameliorating ALD. The application of GSL and favorable GM can be orchestrated beneficially against ALD. However, one of the limitations in this study is that more datasets should be assembled to construct a standard network-based model. It will be the next challenge in the translational medicine.

## Materials and methods

### The acquisition of MEGSL

*Gleditsia sinensis Lam.* (GSL) (**Figure 2A**) was collected from (latitude: 36.176, longitude:128.933) Kyoungsang buk-do, Korea, in September 2022. The GSL was dried at ambient air temperature (22-24 ℃, 10 days), which made it fine powder. The 20 g of GSL powder was soaked with 500ml methyl alcohol (Samchun, Korea, Seoul) for 14 days and repeated twice to obtain greater yield rate. The extract was collected, filtered Whatman filter paper No. 1 (Whatman, Model no. WF1-1850, UK Maidstone), and evaporated with a vacuum evaporator (IKA-RV8, Staufen, Germany) at 40 °C. The obtained amount after evaporating it was 5.72 g (yield rate: 28.6%), which was calculated with below formula: Yield rate (%) = (Evaporated extraction weight/Dried CKF weight) × 100

### GC–MS analysis

GC–MS was adopted to identify the bioactive compounds in GSL, its column was chosen as DB-5 (30 m × 0.25 mm × 0.25 µm) (Agilent, Santa Clara, CA, USA) with reverse phase property. The GC–MS was maintained at 100 °C for 2.1 min, increasing gradually to 300 °C at a rate of 25 °C/min and was kept for 20 min. Injection port temperature was set up at 250 °C and helium flow rate was 1.5 mL/min, and the ionization voltage fixed at 70 eV. The split mode of sample injection was at 10:1 and the Mass Spectra (MS) detection range was between 35 and 900 (m/z). The fragmentation modes of MS were comparative analyzed with the W8N05ST Library MS database (analyzed 10 March 2023).

### The filtration of molecules in GSL

The detected compounds via GC-MS have lipophilic properties, each compound converted into Simplified Molecular Input Line Entry System (SMILES) form via PubChem (https://pubchem.ncbi.nlm.nih.gov/) (accessed 13 March 2023). The molecules were filtered by Lipinski’s rule as drug-likeness selective criterion through SwissADME (http://www.swissadme.ch/) (accessed 13 March 2023). The topological surface area (TPSA) as a representative parameter to identify the cell permeability was considered as a critical factor, its threshold was less than 140 Å^2^ ^[22]^. All parameters’ notations for the molecules have been focused on physicochemical properties to be new agents.

### The extraction of targets related to GSL molecules and ALD

The targets associated with GSL were identified by Similarity Ensemble Approach (SEA) (https://sea16.docking.org/) (accessed on 2 April 2023) and Swiss Target Prediction (STP) (http://www.swisstargetprediction.ch/) (accessed on 2 April 2023), consequentially, selected the intersecting targets between the two databases to achieve more rigorousness. The distinct targets were recognized by Venn diagram plotter. Also, ALD targets were gathered accessing DisGeNet (https://www.disgenet.org/) (accessed on 2 April 2023), and OMIM (https://www.omim.org/) (accessed on 2 April 2023). Ultimately, the overlapping targets obtained between the intersecting targets of GSL and ALD targets were considered as key targets in this study.

### PPI networks and signaling pathway(s)

The final targets mentioned above were extracted by STRING database (https://string-db.org/) (accessed on 5 April 2023), which was described protein protein interaction (PPI) networks to find a key target via R Package. A target with the highest degree value (DV) was defined as the most valuable protein-coding gene in treating ALD. In parallel, a bubble plot based on rich factor (the proportion of target number expressed differentially in each pathway) was constructed to identify the key signaling pathways related to ALD, the information of which was extracted via STRING database (https://string-db.org/) (accessed on 5 April 2023), finalized by R Package.

### The GGSTM networks

The work to merge GSL’s targets, ALD targets, and GM’s targets was realized via GSL or GM-Signaling pathways-Targets-Metabolites (GGSTM) networks construction. The identification of GM metabolite was culled by gutMGene (http://bio-annotation.cn/gutmgene) (accessed on 5 April 2023) with collective resource of GM targets ^[23]^. The description was constructed by R Package. The GM, signaling pathways, targets, and metabolites were represented by nodes based on size of each component. The GM stood for green circle, signaling pathway was symbolized red color, target was illustrated orange color, and metabolite was depicted purple color, all of which were defined as node. The grey line represented the relationships between each node, which was determined as edge. This illustration was performed by R Package.

### The molecular docking test (MDT)

AutoDockTools-1.5.6 was employed to investigate the affinity between key targets and metabolites. For the preparation of key metabolites, each metabolite format was extracted as .sdf file via PubChem (https://pubchem.ncbi.nlm.nih.gov/) (accessed on 7 April 2023), which was converted .sdf file into .pdb file through pymol. The prepared .pdb file was reformatted as .pdbqt file by AutoDockTools-1.5.6. The key target was identified by RCSB Protein Data Bank (PDB), the initial format to be extracted was .pdb file, which was altered into .pdbqt file through AutoDockTools-1.5.6. The AutoDockTools-1.5.6 was established with 4 energy ranges and 8 exhaustiveness to verify the docking score between key targets and molecules. To dock with the conformer (key target-molecule) on the setting boundary, a cubic box was assembled with targets’ active site on 40 Å × 40 Å × 40 Å. The potential centers of the molecule(s) bound to six targets were X: 9.020, Y: 4.548, Z: -19.525; X: 23.285, Y: 12.431, Z: 86.636; X: 90.499, Y: -17.281, Z: -16.848; X: 8.006, Y: -0.459, Z: 23.392; X: 39.265, Y: -18.736, Z: 119.392; and X: 2.075, Y: 31.910, Z: 18.503. The hydrophobic and hydrophilic interactives of the conformers were depicted by LigPlot+ 2.2 (https://www.ebi.ac.uk/thornton-srv/ software/LigPlus/download.html) ^[24]^. The threshold of MDT was determined as -6.0 kcal/mol ^[25]^, and a conformer with the lowest docking score was defined as the most significant regulatory signal in treating ALD.

### The reactivity of key chemical compounds via frontier molecular orbitals (FMO)

Through Lee-Yang-Parr (LYP) correlation functional theory at the 6-31G++ (d,p), the key chemical compounds including positive controls (standard drugs) were completely established by incorporating into GaussView 6.0 and Gaussian 16W software. This optimized setup was to obtain frontier molecular orbitals (FMO) of the lowest unoccupied molecular orbital (LUMO), and highest occupied molecular orbital (HOMO). To assess the chemical value of key chemical compounds, energy gap (eV), hardness (ɳ), softness (S), and electronegativity (χ) were calculated by below formulae.

Energy gap = HOMO – LUMO

ɳ = (LUMO – HOMO)/2

S = 1/ ɳ

χ = – (LUMO – HOMO)/2

The most suitable molecule(s) can be specified with lower energy gap, hardness, and electronegativity in contrast, higher softness ^[26,27]^.

## Abbreviations

AFL: Alcoholic Fatty Liver
AFLD: Alcoholic Fatty Liver Disease
ALD: Alcoholic Liver Disease
DFT: Density Functional Theory
DV: Degree Value
ECM: ExtraCellular Matrix
E_GAP_: Energy gap
FLD: Fatty Liver Disease
FMO: Frontier Molecular Orbital
GC-MS: Gas Chromatography-Mass Spectrometry
GGSTM: GSL or GM-Signaling pathways-Targets-Metabolites
GM: Gut Microbiota
GSL: *Gleditsia sinensis Lam.*
HC: Hepatic Cirrhosis
HCC: HepatoCellular Carcinoma
HOMO: Highest Occupied Molecular Orbital
HS: Hepatic Steatosis
HSCs: Hepatic Stellate Cells
LUMO: Lowest Unoccupied Molecular Orbital
LYP: Lee-Yang-Parr
MDT: Molecular Docking Test
MS: Mass Spectra
MT: Molecular Theory
NP: Network Pharmacology
PPI: Protein Protein Interaction
SEA: Similarity Ensemble Approach
SMILES: Simplified Molecular Input Line Entry System
STP: SwissTargetPrediction
TPSA: Topological Polar Surface Area

## Supplementary Information

Table EV1: The number of 47 chemical constituents from GSL via GC-MS analysis.

Table EV2: The profiled key targets related to ALD via GSL metabolomics.

Table EV3: The degree value (DV) in PPI networks.

Table EV4: The description of key signaling pathways related to ALD.

Table EV5: The binding affinity of key molecules on each target.

## Acknowledgements

Open access publishing facilitated by Hallym University, the Basic Science Research Program through the National Research Foundation of Korea (NRF) funded by the Ministry of Education, Science and Technology, Korea Institute for Advancement of Technology, and Bio Industrial Technology Development Program funded by the Ministry of Trade, Industry and Energy (MOTIE, Korea).

## Author Contributions

DJK and KTS investigated the project. KTS. and KKO designed the study. SBL, SYL, HG, RG, SPS, SMW, and JJJ contributed the data acquisition. KKO, and SJY performed the visualization of the data. KKO, SJY, SBL, and SYL analyzed and interpreted the data. KTS and KKO drafted the manuscript. All authors have read and agreed to the published version of the manuscript.

## Data availability

All data generated or analyzed during this study are included in this published article (and its Supplementary Information files). Additional data related to this study may be requested from the authors.

## Ethics approval and consent to participate

Not applicable.

